# A P300 brain-computer interface with a reduced visual field

**DOI:** 10.1101/2020.09.07.285379

**Authors:** Luiza Kirasirova, Vladimir Bulanov, Alexei Ossadtchi, Alexander Kolsanov, Vasily Pyatin, Mikhail Lebedev

## Abstract

A P300 brain-computer interface (BCI) is a paradigm, where text characters are decoded from visual evoked potentials (VEPs). In a popular implementation, called P300 speller, a subject looks at a display where characters are flashing and selects one character by attending to it. The selection is recognized by the strongest VEP. The speller performs well when cortical responses to target and non-target stimuli are sufficiently different. Although many strategies have been proposed for improving the spelling, a relatively simple one received insufficient attention in the literature: reduction of the visual field to diminish the contribution from non-target stimuli. Previously, this idea was implemented in a single-stimulus switch that issued an urgent command. To tackle this approach further, we ran a pilot experiment where ten subjects first operated a traditional P300 speller and then wore a binocular aperture that confined their sight to the central visual field. Visual field restriction resulted in a reduction of non-target responses in all subjects. Moreover, in four subjects, target-related VEPs became more distinct. We suggest that this approach could speed up BCI operations and reduce user fatigue. Additionally, instead of wearing an aperture, non-targets could be removed algorithmically or with a hybrid interface that utilizes an eye tracker. We further discuss how a P300 speller could be improved by taking advantage of the different physiological properties of the central and peripheral vision. Finally, we suggest that the proposed experimental approach could be used in basic research on the mechanisms of visual processing.

## Introduction

Following the introduction of P300 speller in 1988 by Farwell and Donchin (Farwell and Donchin 1988), many studies have strived to improve this method (Fazel-Rezai, Allison et al. 2012). The speller performance is hindered by the necessity to run many trials to distinguish target and non-target stimuli based on the comparison of the VEPs they evoke. Particularly, non-target items that are adjacent to the target object attract attention and interfere with the decoding performance (Fazel-Rezai 2007, Townsend, LaPallo et al. 2010). Several solutions to this problem have been explored, including using flashes of single items instead of flashing rows and columns (Guger, Daban et al. 2009), rearranging the spatial configuration of the simultaneously flashing stimuli (Townsend, LaPallo et al. 2010), suppressing the stimuli adjacent to targets (Frye, Hauser et al. 2011) or all non-targets (Shishkin, Nikolaev et al. 2011) during the calibration procedure, and optimizing the characteristics of visual stimuli (Salvaris and Sepulveda 2009, Jin, Allison et al. 2012). Yet, all these approaches require a considerable amount of distracting stimuli for accurate spelling, which slows the decoding and causes user fatigue (Boksem, Meijman et al. 2005, Käthner, Wriessnegger et al. 2014).

In this perspective, we discuss a rather straightforward way to reduce the interference from non-target stimuli by the restriction of user sight to only the central visual field. Since the contribution from non-targets is blocked, the response to target stimulus becomes cleaner and easier to detect. Previously, a similar idea was implemented in a single-stimulus BCI for generating an urgent command like braking a neurally-controlled wheelchair (Rebsamen, Guan et al. 2010) or stopping a robot (Fedorova, Shishkin et al. 2014). In these implementations, the speed of operation increased because the decoding was reduced to detecting the presence or absence of the target stimulus. Here we tackled a different approach, where non-targets were effectively removed but the BCI was used for spelling instead of issuing a single command. We have conducted a pilot experiment where visual field reduction was accomplished by wearing a binocular aperture. When looking at the screen through the aperture, subjects were able to perform the same spelling task as they executed with the traditional P300-speller. As intended, the aperture reduced the VEPs caused by non-target stimuli in all subjects. Additionally, responses to the target stimuli became more distinct in 4 out of 10 subjects whereas in the other 6 they were little changed. Thus, the contrast increased between the target and non-target stimuli – an effect that could improve the performance of P300 spellers.

While wearing an aperture was the simplest way to reduce the visual field, alternative approaches could achieve the same goal. Interference from non-tartes could be accomplished algorithmically and/or using a hybrid BCI that incorporates eye tracking (Koo, Nam et al. 2014, Shishkin, Nuzhdin et al. 2016, Hong and Khan 2017). We suggest that the visual field reduction method could be useful for both clinical applications and studies of visual processing and attention.

## Speller settings with and without the aperture

The pilot experiment was conducted as a part of our ongoing experiments on a P300 BCI of a traditional design. Ten healthy subjects first performed a traditional P300 task and then switched to wearing a binocular aperture that restricted their sight to the central visual field (Figure 1).

**Figure 1.**
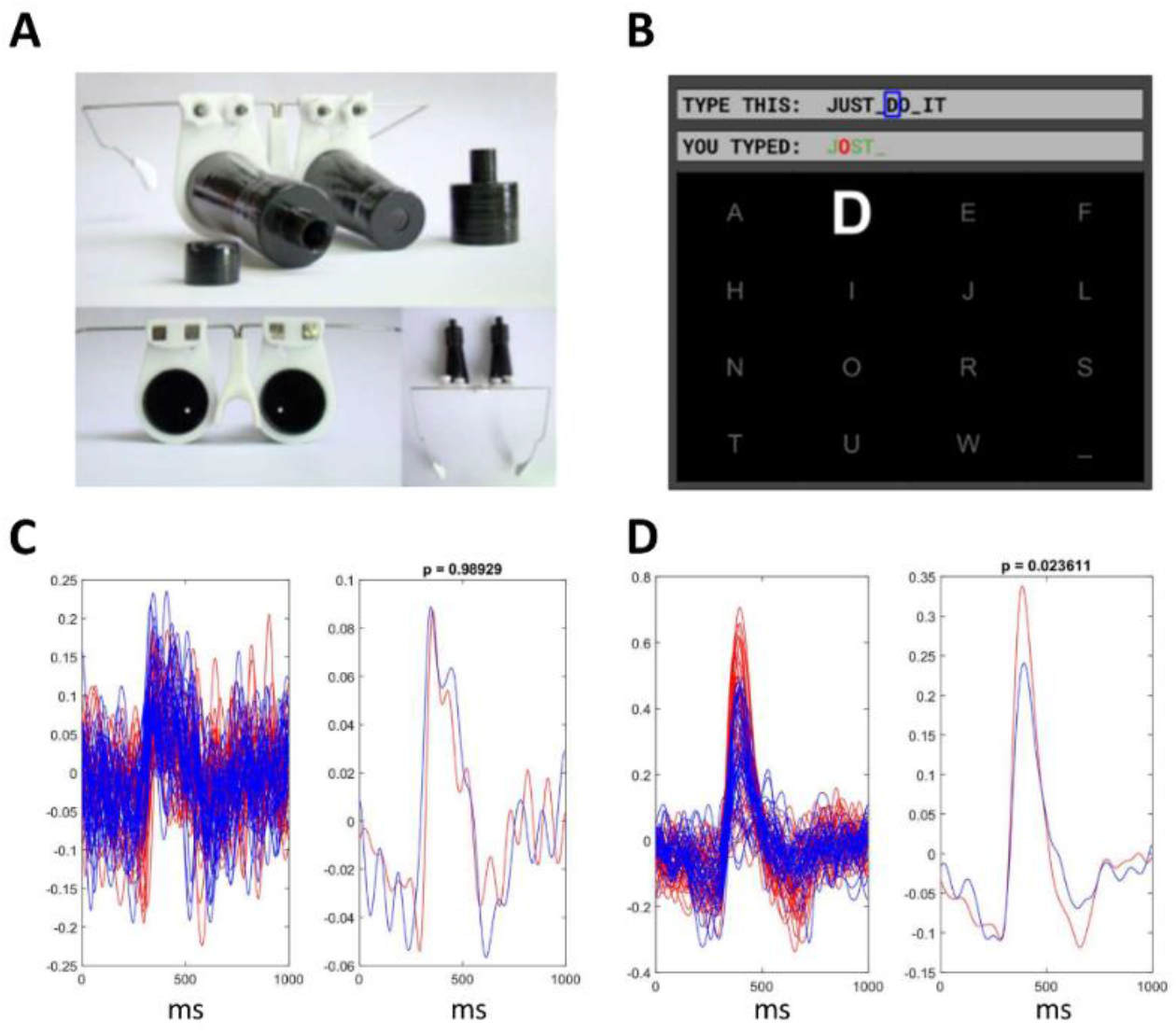
Experimental setup and example VEPs in response to target stimuli. A: Aperture headset. B: Computer screen with text characters in a 4 by 4 arrangement. C: VEPs in a subject with the response unchanged during wearing the aperture. Red lines correspond to the aperture condition, blue lines represent the no-aperture condition. Average VEPs for 10 flashes (i.e. one character spelled) are shown on the left, and average VEPs for all flashes (i.e. a complete phrase spelled) are shown on the right. D: VEPs in a subject with an enhanced response during the aperture condition.

The experimental procedure was approved by the research ethics committee of Samara State Medical University (protocol #204, December 11, 2019). All subjects gave informed consents to their participation in the study. Ten healthy subjects, all males, aged 19±0.0, right-handed, with right dominant eye determined with Miles test (Miles 1929), with visual acuity of 1.04±0.18 and 1.09±0.18 for the left and right eyes, respectively, measured with Huvitz CCP-3100 projector (HUVITZ Co., Ltd, S.Korea) using Cyrillic letters.

The experiments were conducted in a half-dimmed quiet room. Participants sat in front of a 40×70 cm computer monitor. The distance from the eyes to the screen was 80 cm. The BCI system incorporated an NVX-36 amplifier, NeoRec software (MKS, Russia), OpenViBE software (Renard, Lotte et al. 2010), VIBRAINT software, and an aperture headset (IT Universe Ltd). EEG channels P4, PZ, P4 were recorded according to the international “10-20” system. The left earlobe was used as the reference channel and the right earlobe as the ground. We used a textile cap with gel-based Ag/AgCl wired electrodes MCScap-E (MKS, Russia). The sampling rate of the NVX-36 amplifier amplifier was set to 250 Hz. The EEG signal from the NVX-36 amplifier was received by OpenViBE Acquisition software and then transmitted to VIBRAINT software for processing. In the experiments with the aperture, the aperture headset was fixed to the head like regular glasses (Figure 1A). The aperture tubes were coated with an anti-glare spraying on the inside. The aperture opening was 5 mm in diameter. The distance from the openings to the eyes was 12 cm. Subject could see only one character at a time when wearing the aperture.

Server software and then transmitted to VIBRAINT software for processing. In the experiments with the aperture, the aperture headset was fixed to the head like regular glasses (Figure 1A). The aperture tubes were coated with an anti-glare spraying on the inside. The aperture opening was 5 mm in diameter. The distance from the openings to the eyes was 12 cm. Subject could see only one character at a time when wearing the aperture.

In our implementation of the P300 speller, 16 characters were displayed on a computer screen in a 4 by 4 arrangement (Figure 1B). The characters flashed randomly, one at a time. When flashing, the character doubled in size (i.e. linear dimensions), and its brightness increased by 80%. Flashes occurred every 110 ms (i.e. 9.1 Hz). Flash duration was 60 ms. Subjects used the BCI to generate a ten-character phrase (“JUST DO IT”), first with the traditional P300 speller and then while wearing the aperture. Prior to running each speller tasks, a calibration session was conducted, where data were collected needed for fitting an SVM-based classifier. The classifier was implemented in VIBRAINT software. Participants were instructed to look at the target character and avoid making head movements. For every character to be spelled, each screen item flashed 10 times. Next, the classifier predicted the target stimulus based on these data. The classification result was displayed on the screen. For example, in the experimental session shown in Figure 1B, the classifier correctly determined “J”, “S”, “T” and space, and confused “U” with “O”.

## Aperture effect on VEPs

The comparison of VEPs across the experimental conditions (with and without the aperture) was conducted offline. EEG signals were preprocessed for this analysis by applying a bandpass filtered in the range 1.0-15.0 Hz with a 4th order Butterworth IIR zero-phase filter. Additionally, EEGs on each channel were normalized to bring the amplitude to the [-1, 1] range. We used a PZ channel to assess VEPs in different conditions.

Wearing the aperture resulted in a significant change of VEP in response to target stimulus (p<0.05 Wilcoxon rank-sum test for response amplitude) in 4 subjects out of 10, and no change in the rest of the subjects. Figure 1C shows the responses to target stimuli for a subject with no change in target-stimulus VEP occurring due to wearing the aperture. Blue lines correspond to regular P300 runs, and red lines correspond to the aperture condition. Traces shown in the left panel represent averages for 10 target flashes, and traces in the right panel represent averages for all flashes. In this case, VEPs are very similar in both conditions (Figure1C, right panel) and there is no statistically significant difference. Figure 1D shows the results for a subject with an approximately 40% increase in VEP amplitude when wearing the aperture; this effect is statistically significant.

Figure 2 shows VEPs (averages for all flashes) for both target stimuli (left parts of the panels) and non-target stimuli (right parts). Figure 2A and Figure 2C represent the same subjects as in Figure 1C and Figure 1D, respectively. In these examples, there is no significant change in the amplitude of VEP to target stimulus in two subjects (Figure 2A,B), and the response is stronger in the aperture condition in four subjects (Figure 2C-F).

**Figure 2.**
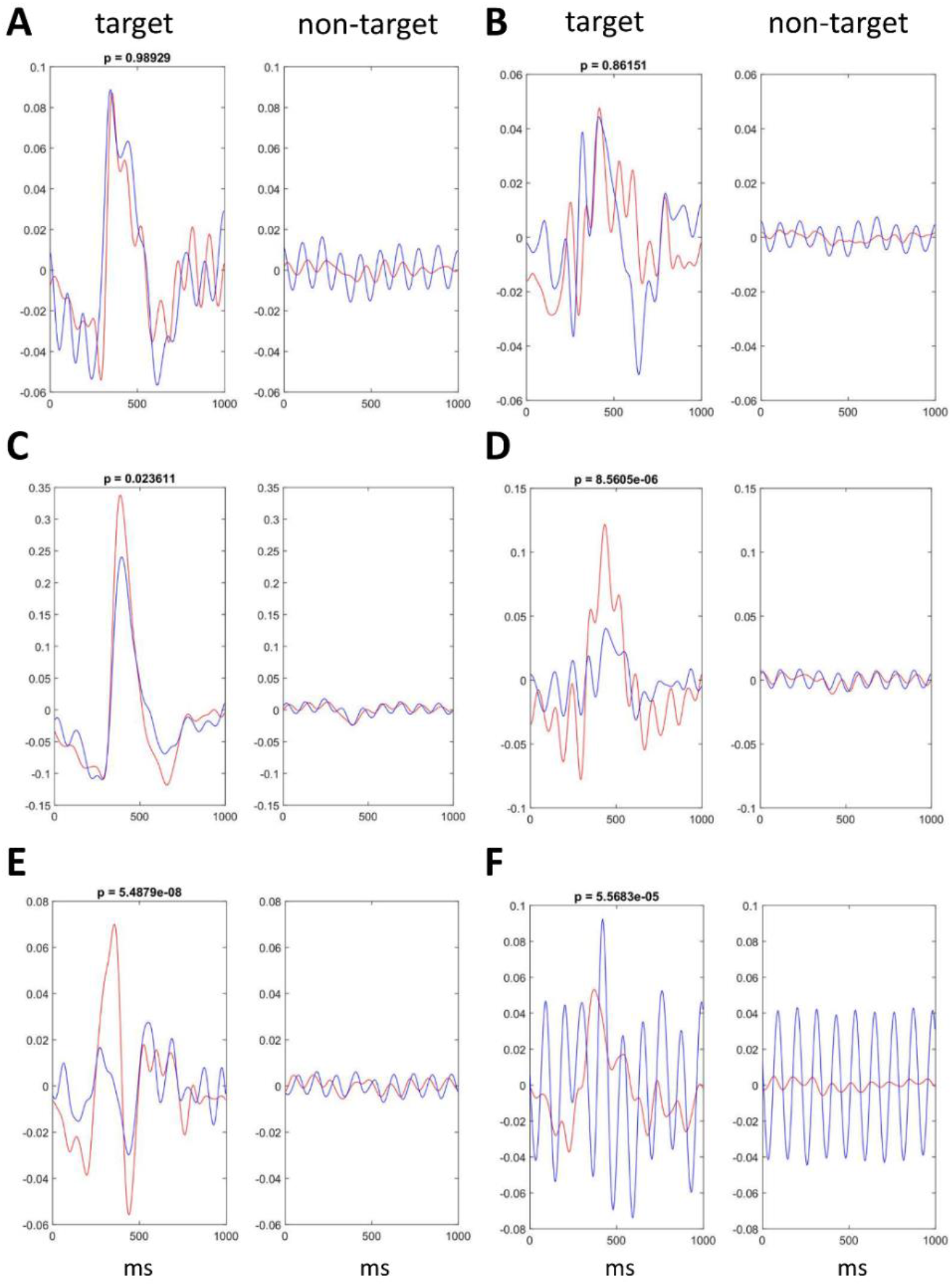
VEPs for target (left panels) and non-target (right panels) stimuli. Traces represent averages for all flashes. Blue lines are the responses for the regular P300 paradigm, and red lines are the responses for the aperture condition, where the visual field was reduced to central vision. Panels A and C show the data for the same subjects as in Figure 1C and Figure 1D, respectively. There is no change in target-stimulus response in panels A and B whereas the other panels contain cases with such a change. In all cases, the response to non-targets decreased when subjects wore the aperture.

Moreover, in all cases there is a decrease in the response to non-target stimulus in the aperture condition. A particularly strong decrease in the non-target response is shown in Figure 2F. In this subject, cortical response was strong for both target and non-target stimuli during the regular P300-BCI runs, and the responses to non-target stimuli were abolished by wearing the aperture. Overall, the aperture worked as designed. It diminished the responses to the interfering items, and in several subjects, target-stimulus VEP increased. These both effects enhanced the contrast between the target and non-target responses, which suggests that reducing the visual field to central vision could be useful to improve the performance of P300 speller.

## Discussion

Our pilot experiment showed that subjects could operate a P300 speller while wearing an aperture that restricted their visual field to central vision. With this method, the interference from non-target stimuli was reduced, as evident from the VEP data. Moreover, in 4 subjects out of 10, we observed changes in cortical responses to target stimuli: in 3 subjects that was an increase in VEP amplitude and in one subject that was a conversion of a noisy mixture of target and non-target responses into a clean response pattern. These preliminary observations support our hypothesis that removal of non-target stimuli could help distinguish targets from non-targets and even remove the need to present non-targets in the VEP-based speller paradigm.

Our implementation of the reduced visual field idea could be improved in the future. Looking at the screen through a fixed-sized aperture is uncomfortable, and this problem could be addressed using a variable-size aperture that narrows the visual field only when needed. Additionally, the aperture could be applied to one eye only and the subject could close the other eye when focusing on the target. Furthermore, the same effect could be achieved with an algorithm that reduces the flicker of non-targets as VEP data accumulates, and eventually stops it. A hybrid BCI that incorporates eye tracking (Koo, Nam et al. 2014, Shishkin, Nuzhdin et al. 2016, Hong and Khan 2017) could be also used for this purpose. Indeed, once the fixation point is detected by an eye-tracker, flashes of non-target items could be suppressed. With such a method, the BCI could be controlled without producing head movements, which were necessary in our aperture-based implementation.

Regardless of the technical details of how this approach could be implemented, we see several advantages in a speller where only the central visual field is used for stimulus delivery. The first advantage is that distractors do not interfere with selective spatial attention (Clark and Hillyard 1996, Hillyard and Anllo-Vento 1998, Luck, Woodman et al. 2000, Ordikhani-Seyedlar, Lebedev et al. 2016). This consideration is particularly important when using the speller in patients who may have impaired attention in addition to the other neurological conditions (Vieregge, Wauschkuhn et al. 1999, Phukan, Pender et al. 2007, Nazhvani, Boostani et al. 2013, Yadav, Thiagarajan et al. 2014). The second advantage is the reduction of visual fatigue, which is a common problem for BCI spellers (Boksem, Meijman et al. 2005, Käthner, Wriessnegger et al. 2014). Finally, the third obvious advantage is that, when the number of non-target stimuli is reduced, the speller works faster.

One could argue that, since we are suggesting that non-target stimuli could be eliminated completely, the same could be achieved with an eye tracker and/or head-position tracker: once the gaze angle is reliably detected, spelling could be performed without the need to use a BCI. While this solution could be practical in many cases, recording brain activity is still very useful (Shishkin, Nuzhdin et al. 2016) because it allows to better assess the subject’s intention, level of attention, and the brain state. Additionally, eye/head tracking is not always possible. We therefore suggest that the most versatile approach would be using as many control signals as possible and selecting the appropriate ones for each concrete task (Hong and Khan 2017).

Even if the visual field reduction does not happen to be practical in some cases, it could be still useful for the assessment of an individual subject’s ability to control a BCI. For example, in our study, target-stimulus VEPs were unchanged for 6 subjects wearing the aperture, whereas they were affected in 4 other subjects, which suggests that the latter subjects focused better on the target in the absence of distractors. In one of these 4 subjects, responses to non-targets were extraordinarily strong in the regular P300-BCI sessions possibly because of the strong engagement of exogenous attention by the distractors. The removal of distractors dramatically improved the cleanness of the response to the target stimulus, so at least in this subject wearing the aperture had a major positive effect. With more subjects tested in the future and this kind of paradigm extended to patients, we expect that more useful information will be obtained regarding the usefulness of our approach in different individuals.

Even the simple aperture test implemented here could serve as useful control that is quickly run to assess the VEPs with and without distracting stimuli. (Such a control does not require changing the basic P300 paradigm, so it is easy to implement.)

In addition to bringing improvements to the BCI speller and providing helpful controls, the paradigm proposed here could be useful as a tool in research on brain mechanisms of visual processing and attention. Thus, it is well known that central and peripheral visual fields have different functional properties (Finlay 1982, Neville and Lawson 1987, Berencsi, Ishihara et al. 2005), but this topic has not been sufficiently studied with the BCI approach. Given that BCI research contributes to basic science (Nicolelis and Lebedev 2009), we expect more fundamental insights on the brain mechanisms with the proposed approach.

## Acknowledgement

This manuscript has been released as a pre-print at biorxiv.org (Kirasirova, Bulanov et al. 2020). ML proposed the idea, LK, VB, AO, AK, VP and ML designed the experiments. LK, VB and VP conducted experiments, LK, VB, AO, and ML analyzed data, LK, VB, AK, VP and ML wrote the manuscript. Author VB was employed by the company IT Universe Ltd, Samara, Russia. The remaining authors declare that the research was conducted in the absence of any commercial or financial relationships that could be construed as a potential conflict of interest. We thank Ilia Borishchev, Maxim Averkiev, Dmitry Kuchkin, Oleg Mukhin, Yuri Potantsev, Tatiana Zubkova (IT Universe Ltd) for their contribution to software and equipment development and Victoria Germanova for the help with testing vision of the subjects. The study was funded by RFBR, project number No 19-315-90120.

## Notes

### Competing Interest Statement

The authors have declared no competing interest.

### Summary of Updates

Several references added; text slightly revised.

